# Evaluating fast maximum likelihood-based phylogenetic programs using empirical phylogenomic data sets

**DOI:** 10.1101/142323

**Authors:** Xiaofan Zhou, Xingxing Shen, Chris Todd Hittinger, Antonis Rokas

**Author notes:** Correspondence: Antonis Rokas.

## Abstract

Phylogenetics has witnessed dramatic increases in the sizes of data matrices assembled to resolve branches of the tree of life, motivating the development of programs for fast, yet accurate, inference. For example, several different fast programs have been developed in the very popular maximum likelihood framework, including RAxML/ExaML, PhyML, IQ-TREE, and FastTree. Although these four programs are widely used, a systematic evaluation and comparison of their performance using empirical genome-scale data matrices has so far been lacking. To address this question, we evaluated these four programs on 19 empirical phylogenomic data sets from diverse animal, plant, and fungal lineages with respect to likelihood maximization, tree topology, and computational speed. For single-gene tree inference, we found that the more exhaustive and slower strategies (ten searches per alignment) outperformed faster strategies (one tree search per alignment) using RAxML, PhyML, or IQ-TREE. Interestingly, single-gene trees inferred by the three programs yielded comparable coalescent-based species tree estimations. For concatenation–based species tree inference, IQ-TREE consistently achieved the best-observed likelihoods for all data sets, and RAxML/ExaML was a close second. In contrast, PhyML often failed to complete concatenation-based analyses, whereas FastTree was the fastest but generated lower likelihood values and more dissimilar tree topologies in both types of analyses. Finally, data matrix properties, such as the number of taxa and the strength of phylogenetic signal, sometimes substantially influenced the relative performance of the programs. Our results provide real-world gene and species tree phylogenetic inference benchmarks to inform the design and execution of large-scale phylogenomic data analyses.

## Introduction

Phylogenetic analysis – that is, the identification of the tree best representing the evolutionary history of the underlying data – is of fundamental importance to many biological disciplines, including but not limited to systematics, molecular evolution, and comparative genomics (Felsenstein 2003; Xia 2013; Hamilton 2014; Yang 2014). However, finding the best tree is an exceptionally difficult task because evaluation of each tree requires a considerable amount of calculations (Bryant et al. 2005) as well as because the number of candidate strictly bifurcating trees grows very rapidly with the number of sequences (Felsenstein 1978) – for example, there are ∼8 × 10^21^ possible rooted topologies for a set of 20 taxa. Therefore, fast programs that employ heuristic algorithms that can efficiently infer the best tree (or nearly as good alternatives) are of pivotal importance to phylogenetic analysis. This is evident by the success of the Neighbour-Joining (NJ) method, a distance-based clustering (instead of tree searching) algorithm (Saitou and Nei 1987) that is the most highly cited phylogenetic method (Van Noorden et al. 2014). NJ and its variants (e.g. BIONJ which takes the variance of distance estimation into consideration) (Gascuel 1997; Bruno et al. 2000) were among the few available options for analyzing large data sets until the 2000s, and are still widely used today to quickly produce good starting points for more sophisticated methods (e.g. Guindon et al. 2010; Nguyen et al. 2015).

It is now generally accepted that statistical methods, such as maximum likelihood (ML) (Felsenstein 1981), produce more reliable results than distance and parsimony methods (Yang and Rannala 2012; Whelan and Morrison 2017). However, ML-based methods are also computationally more expensive, necessitating the use of heuristic search algorithms for searching the enormity of tree space (Chor and Tuller 2005). Heuristic search algorithms typically adopt iterative, “hill-climbing” optimization techniques that involve three steps: (1) generate a quick starting tree (e.g. BIONJ tree, stepwise-addition parsimony tree, etc.); (2) modify the tree using certain topological rearrangement rules and evaluate the resultant trees under the ML criterion; and (3) replace the starting tree and repeat step 2 if the rearrangements identify a better tree, or otherwise terminate the search. The most common rearrangement algorithms for step 2 are Nearest-Neighbor-Interchange (NNI), where the four subtrees connected by a given internal branch are re-arranged to form two new, alternative topologies (Robinson 1971), and Subtree-Pruning-and-Regrafting (SPR), in which a given subtree is detached from the full tree and re-inserted onto each of the remaining branches (Swofford et al. 1996). SPR is more expansive in searching tree space than NNI since it can evaluate many more trees from one initial topology, but it is also much slower because of the extra tree evaluations.

Four of the most popular fast ML-based phylogenetic programs that differ in their choices or implementations of rearrangement algorithms are PhyML (Guindon and Gascuel 2003; Guindon et al. 2010), RAxML / ExaML (Stamatakis 2014; Kozlov et al. 2015), FastTree (Price et al. 2010), and IQ-TREE (Nguyen et al. 2015). First introduced in the early 2000s, PhyML has been one of the most widely used programs for ML-based phylogenetic inference (Guindon and Gascuel 2003). The original algorithm was based solely on NNI and achieved comparable performance as other contemporary ML methods but with much lower computational costs. The latest version of PhyML (version 20160530) performs hill-climbing tree searches using SPR rearrangements in early stages and NNI rearrangements in later stages of the tree search (Guindon et al. 2010). Specifically, during the SPR-based search, candidate re-grafting positions are first filtered based on parsimony scores; the most parsimonious ones are then subject to approximate ML evaluation where branch-lengths are only re-optimized at the branches adjacent to the pruning and re-grafting positions. To accelerate the tree search, the best “up-hill” SPR move for each subtree is accepted immediately, potentially leading to the simultaneous application of multiple SPRs in one round. Once the search has converged to a single topology, the resultant tree is further optimized by NNI-based hill-climbing. Similar to the SPR stage, PhyML evaluates candidate NNIs only approximately by re-optimizing the five relevant branches, and may apply multiple NNI moves simultaneously at each round. The addition of the SPR algorithm in PhyML has significantly improved its accuracy, although at the cost of longer runtimes (Guindon et al. 2010).

RAxML is another widely used program for fast estimation of ML trees (Stamatakis 2006, 2014). The latest version (8.2.11) implements the standard SPR-based hill-climbing algorithm and employs important heuristics to reduce the amount of unpromising SPR candidates, including: 1) candidate re-grafting positions are limited to only those within a certain distance from the pruning position (known as the “lazy subtree rearrangement”) (Stamatakis et al. 2005); and 2) if the re-grafting to a candidate position results in a substantially worse likelihood value, all branches further away from that point will be ignored (Stamatakis et al. 2007). As in PhyML, the approaches of approximate pre-scoring of SPR candidates and simultaneous SPRs are also used by RAxML to speed up the analysis (Stamatakis et al. 2005). In addition to RAxML, its sister program ExaML is specifically engineered for large concatenated data sets (Kozlov et al. 2015); it achieves greatly enhanced parallel efficiency through a novel balance load algorithm and parallel I/O optimization. As RAxML has exhibited excellent performance in both accuracy and speed (Stamatakis 2006), it is considered by many to be the state-of-the-art ML fast phylogenetic program.

Although both PhyML and RAxML represent great advances in developing fast and accurate phylogenetic programs, efforts aimed at improving the speed of ML tree estimation continue. For example, the recently developed FastTree program can be orders of magnitude faster than either PhyML or RAxML / ExaML (Price et al. 2010). FastTree (latest version 2.1.10) first constructs an approximate NJ starting tree which is then improved under the minimum evolution criterion using both NNI and SPR rearrangements, followed by ML-based NNI rearrangements to search for the final tree. With computational efficiency at the very heart of its design, FastTree makes heavy use of heuristics at all stages to limit the numbers of tree searches and likelihood optimizations. As a tradeoff, FastTree generates less accurate tree estimates than SPR-based ML methods (Price et al. 2010). The substantial edge of the FastTree program in speed has made it very popular, particularly in analyses of very large phylogenomic data matrices.

An important weakness of pure hill-climbing methods is that they can be easily trapped in local optima. The IQ-TREE program, the most recent of the four fast ML-based phylogenetic programs, was developed aiming to overcome this local optimum problem through the use of stochastic techniques (Nguyen et al. 2015). Specifically, IQ-TREE (latest version 1.5.5) generates multiple starting trees instead of one and subsequently maintains a pool of candidate trees during the entire analysis. The tree inference proceeds in an iterative manner; at every iteration, IQ-TREE selects a candidate tree randomly from the pool, applies stochastic perturbations (e.g. random NNI moves) onto the tree, and then uses the modified tree to initiate a NNI-based hill-climbing tree search. If a better tree is found, the worst tree in the current pool is replaced and the analysis continues; otherwise, the iteration is considered unsuccessful and the analysis terminates after a certain number of unsuccessful iterations. IQ-TREE takes advantage of successful preexisting heuristics (e.g. simultaneous NNIs [Guindon and Gascuel 2003]) and a highly-optimized implementation of likelihood functions (Flouri et al. 2015) for better computational efficiency.

These four programs offer different tradeoffs between the extent of tree space searched and speed in fast phylogenetic inference, and they may exhibit different behaviors toward diverse phylogenomic data sets whose properties (e.g. taxon number and gene number) and evolutionary characteristics (e.g. age of lineage, taxonomic range, and evolutionary rate) vary. Therefore, a good understanding of their relative performance across diverse empirical phylogenomic data matrices is critical to the success of phylogenetic inference when computational resources are limited. This is particularly relevant for large-scale studies using data matrices of ever-increasing data volumes and complexities. So far, these four programs have only been evaluated using simulated data (Guindon et al. 2010; Price et al. 2010; Liu et al. 2011), which might not well approximate real data, and relatively small empirical data sets containing ∼10 to ∼200 gene alignments (Guindon et al. 2010; Price et al. 2010; Liu et al. 2011; Money and Whelan 2012; Nguyen et al. 2015; Chernomor et al. 2016), which might lack generality. In these studies, RAxML and PhyML showed largely similar performance in identifying trees of higher likelihood scores (Guindon et al. 2010; Money and Whelan 2012), while IQ-TREE exhibited improved efficiency compared to both RAxML and PhyML (Nguyen et al. 2015; Chernomor et al. 2016). On the other hand, FastTree was found to be much faster than RAxML and PhyML but reported lower likelihood scores for data sets with both small and large numbers of sequences (Guindon et al. 2010; Price et al. 2010; Liu et al. 2011). However, it remains unclear if these patterns would hold for large empirical data sets and for species tree estimation based on genome-scale data.

To comprehensively evaluate the four fast ML-based phylogenetic programs (table 1), we used a large collection of 19 empirical phylogenomic data sets representing a wide range of properties, including data types (both DNA and protein data), numbers of taxa (up to 200) and genes (up to 14,446), and taxonomic range for diverse animal, plant, and fungal lineages (table 2; for details on the source of each data set, see supplementary table S1). For each of these data sets, we compared the performance of all programs for single-gene tree inference and, for coalescent-based and concatenation-based species tree inference, the two major current approaches to inferring species phylogenies from phylogenomic data (Liu et al. 2015). In the coalescent approach, the species tree is estimated by considering all individually inferred single-gene trees using coalescent methods that take into account that the histories of genes may differ from those of species due to incomplete lineage sorting (fig. 1A), whereas in the concatenation approach, the species tree is estimated from the supermatrix derived by concatenating all single-gene alignments (fig. 1B).

**Table 1.** Overview of the four fast ML-based phylogenetic programs evaluated in this study.

| Programs | Optimality<br>criterion | Starting tree | Topological<br>moves | Supported models |  | Partitioned<br>analysis |
| --- | --- | --- | --- | --- | --- | --- |
|  |  |  |  | AA | DNA |  |
| RAxML<br>v8.2.0<br>(ExaML<br>v3.0.17) | ML | Parsimony /<br>random /<br>custom | SPR | Common<br>and custom<br>models | JC69, K80,<br>HKY85,<br>GTR | Y |
| PhyML<br>v20160530 | ML | Parsimony /<br>random /<br>custom | Interleaved<br>NNI and<br>SPR | Common<br>and custom<br>models | Common<br>and custom<br>models | Y |
| IQ-TREE<br>v1.4.2 | ML | BIONJ and<br>multiple<br>parsimony /<br>random /<br>custom | NNI and<br>stochastic<br>perturbation | Common<br>and custom<br>models | Common<br>and custom<br>models | Y |
| FastTree<br>v2.1.9 | ML | Heuristic<br>NJ | NNI and<br>SPR (ME)<br>followed by<br>NNI (ML) | JTT, WAG,<br>LG | JC69, GTR | N |

Note: ML – maximum likelihood; ME – minimum evolution; NJ – neighbor joining; NNI – nearest neighbor interchange; SPR – subtree pruning and re-grafting.

**Figure 1.**
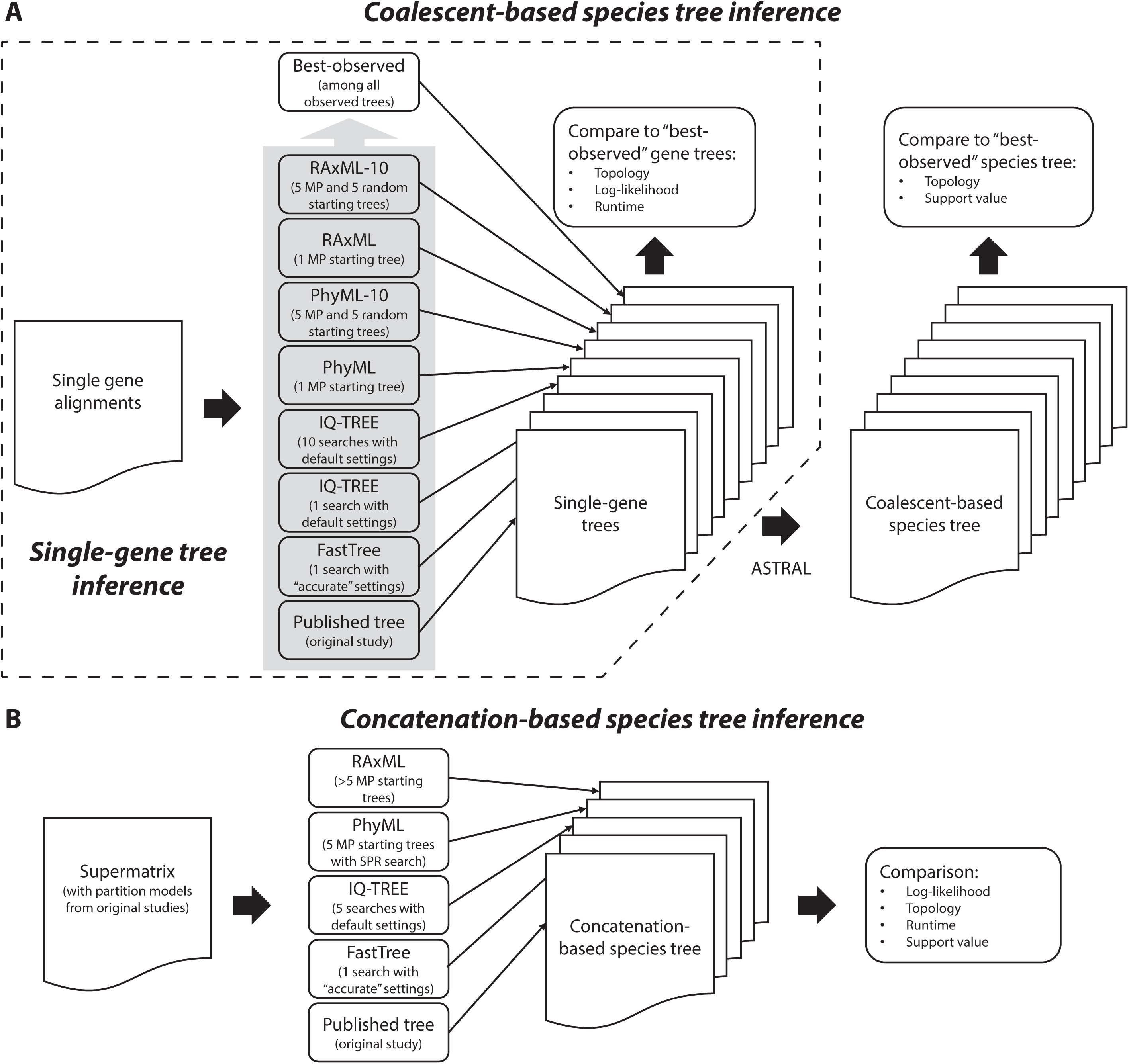
Schematics of the (A) single-gene tree inference test as well as the coalescent-based and (B) concatenation-based species tree inference tests used to evaluate the performance of fast phylogenetic programs in phylogenomic analysis.

**Table 2.** Overview of the 19 phylogenomic data sets included in this study.

| Study | Data set* |  | Genes | Taxa | Taxonomic group | Data type |
| --- | --- | --- | --- | --- | --- | --- |
|  | AA | DNA |  |  |  |  |
| (Nagy et al. 2014) | NagyA1 |  | 594 | 60 | Fungi | Genome |
| (Misof et al. 2014) | MisoA2 |  |  |  |  |  |
|  |  | MisoD2a<sup>1,2</sup> | 1,478 | 144 | Insects | Transcriptome |
|  |  | MisoD2b<sup>2</sup> |  |  |  |  |
| (Wickett et al. 2014) | WickA3 |  |  |  |  |  |
|  |  | WickD3a<sup>2</sup> | 844 | 103 | Land plants | Transcriptome |
|  |  | WickD3b<sup>3</sup> |  |  |  |  |
| (Chen et al. 2015) | ChenA4 |  | 4,682 | 58 | Vertebrates | Transcriptome |
| (Struck et al. 2015) | StruA5 |  | 679 | 100 | Worms | Transcriptome |
| (Borowiec et al. 2015) | BoroA6 |  | 1,080 | 36 | Metazoans | Genome |
| (Whelan et al. 2015) | WhelA7 |  | 210 | 70 | Metazoans | Transcriptome |
| (Yang et al. 2015) | YangA8 |  | 1,122 | 95 | Caryophyllales | Transcriptome |
| (Shen et al. 2016b) | ShenA9 |  | 1,233 | 96 | Yeasts | Genome |
| (Song et al. 2012) |  | SongD1 | 424 | 37 | Mammals | Genome |
| (Xi et al. 2014) |  | XiD4 | 310 | 46 | Flowering plants | Transcriptome |
| (Jarvis et al. 2014) |  | JarvD5a<sup>4</sup> | 14,446 | 48 | Birds | Genome |
|  |  | JarvD5b<sup>4</sup> | 2,022 | 48 |  |  |
| (Prum et al. 2015) |  | PrumD6 | 259 | 200 | Birds | Target |

|  |  |  |  |  | Enrichment |
| --- | --- | --- | --- | --- | --- |
| (Tarver et al. 2016) | TarvD7 | 11,178 | 36 | Mammals | Genome |

\*Data sets are named using the first four letters of the first author's last name from the study the data set was generated, followed by the letter A (for amino acid) or D (for DNA), followed by a unique numeric or alphanumeric identifier.
<sup>1</sup>Data set MisoD2a does not have a corresponding supermatrix from the original study;
<sup>2</sup>DNA data sets MisoD2a and WickD3a include the codon-based alignments corresponding to the amino acid alignments in data sets MisoA2 and WickA3, respectively;
<sup>3</sup>DNA data sets MisoD2b and WickD3b include the full codon-based alignments corresponding to the amino acid alignments in data sets Miso2 and Wick3, respectively, with the third codon positions removed;
<sup>4</sup>Data set JarvD5b were derived from data set JarvD5a through statistical binning (Mirarab et al. 2014), and the two data sets correspond to the same supermatrix.

In single-gene tree estimation, we found that, although the more comprehensive analysis strategy (ten searches per alignment using RAxML, PhyML, or IQ-TREE) performed considerably better than fast strategies (one tree search per alignment using the same programs), all produced results of comparable quality when the inferred gene trees were used for coalescent-based species tree inference. The impact of tree search numbers and starting tree types on the efficiency of single-gene alignment analysis was also investigated. For the concatenation-based species tree inference, we found that, in some cases, IQ-TREE recovered trees with higher likelihood scores than RAxML/ExaML, although both showed the best performance for most data sets. Importantly, IQ-TREE exhibited comparable or better speed in both coalescent-based and concatenation-based species tree inference compared with RAxML/ExaML. In contrast, FastTree produced significantly worse single-gene and species trees than the other three programs even when allowed to run multiple times, whereas PhyML did not scale well to supermatrices because the concatenation-based species tree inferences failed to complete for multiple data sets. Overall, our benchmarking of the four fast ML-based phylogenetic programs against 19 state-of-the-art data matrices is highly informative for the design of efficient data analysis strategies in phylogenomic studies.

## Results and Discussion

### A comprehensive collection of empirical data

For a comprehensive evaluation of the four fast ML-based phylogenetic programs, we retrieved 19 data sets from 14 recently published phylogenomic studies (table 2; see supplementary table S1 for detailed sources of each data set), representing a wide range of characteristics: 1) they include both amino acid and nucleotide data sets (nine and ten, respectively); 2) they contain either moderate numbers of taxa (e.g. PrumD6, 200 taxa and 259 genes [Prum et al. 2015]), large numbers of genes (e.g. JarvD5a, 48 taxa and 14,448 genes [Jarvis et al. 2014]), or both (e.g. MisoA2, 144 taxa and 1,478 genes [Misof et al. 2014]); 3) they cover three major taxonomic groups (i.e. animals, plants, and fungi) and various depths within each group (e.g. data sets SongD1 [Song et al. 2012], ChenA4 [Chen et al. 2015], and WhelA6 [Whelan et al. 2015] cover mammals, vertebrates, and metazoans, respectively); and 4) they consist of sequence data derived from different technologies (e.g. some data sets were built entirely on whole genome sequences [Song et al. 2012; Jarvis et al. 2014; Shen et al. 2016b; Tarver et al. 2016], while some others contained mostly transcriptome sequencing data [Misof et al. 2014; Wickett et al. 2014; Yang et al. 2015]). In addition, these data sets were assembled and curated in state-of-the-art phylogenomic studies and thus are of high quality. Therefore, these data sets are well suited for benchmarking the performance of fast phylogenetic programs in the context of phylogenomics.

### Performance Test I: Single-gene tree inference

In the first test, we examined the performance of four fast ML-based phylogenetic programs (i.e. RAxML, PhyML, IQ-TREE, and FastTree) in inferring single-gene trees (fig. 1A). We designed seven strategies, including four basic strategies in which each program was used to infer each gene tree from a single starting tree (these were named *RAxML*, *PhyML*, *IQ-TREE*, and *FastTree*), as well as three more comprehensive strategies in which each of RAxML, PhyML, and IQ-TREE was used to infer each gene tree from ten replicates (these were named *RAxML-10*, *PhyML-10*, and *IQ-TREE-10*). In both *RAxML-10* and *PhyML-10*, five of the starting trees were obtained via parsimony (including the ones used in the *RAxML* and *PhyML* strategies, respectively) and the other five were random starting trees. On the other hand, *IQ-TREE-10* consists of ten independent IQ-TREE searches, including the one performed in *IQ-TREE*.

The seven strategies were compared for the likelihood scores and topologies of their single-gene tree inferences, as well as for their computational speeds. Since the true evolutionary histories are unknown for the empirical data used here, we identified the tree with the highest likelihood score for each alignment (hereafter referred to as the “best-observed” tree) among trees inferred by the seven strategies and the trees reported in previous studies, if available. These “best-observed” trees were used as the reference in the comparisons of likelihood score and topology.

#### Likelihood score maximization

We first examined the performance of the seven strategies in likelihood score maximization on single-gene alignments (supplementary table S2) by calculating the frequencies with which each of the seven strategies had the highest score (fig. 2). Overall, *IQ-TREE-10* and *RAxML-10* had the highest frequencies of finding the highest likelihood scores (80.17% and 75.99%, respectively) and reported the highest likelihood scores more frequently than the other strategies in all data sets except for JarvD5b, highlighting the benefit of using multiple starting trees. Importantly, the performances of *IQ-TREE-10* and *RAxML-10* varied substantially among data sets; whereas the two strategies performed very similarly on several data sets (e.g. NagyA1 and SongD1), in others *RAxML-10* outperformed *IQ-TREE-10* by large margins (e.g. MisoA2, MisoD2a, and MisoD2b), or vice versa (e.g. JarvD5b).

**Figure 2.**
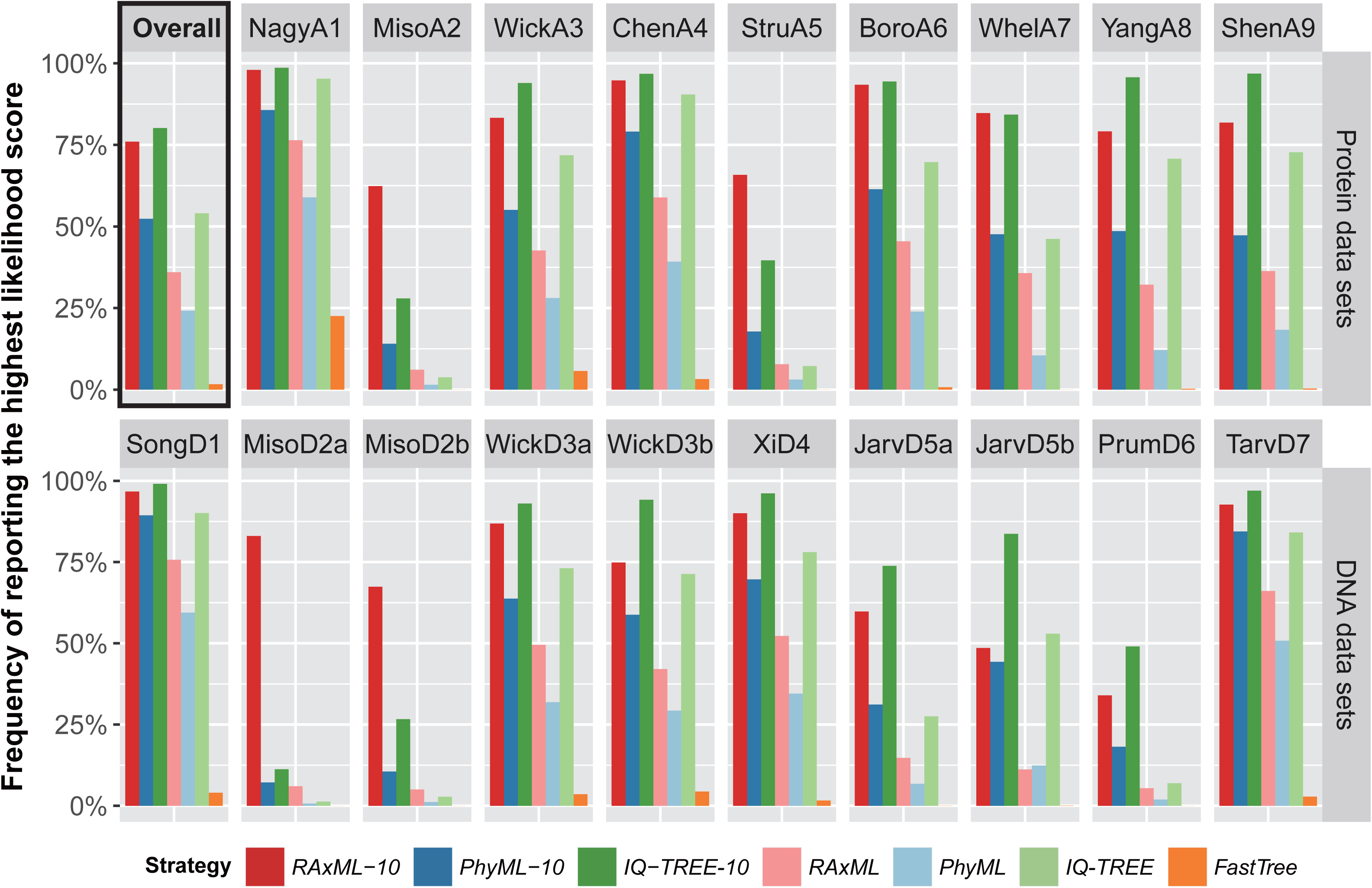
Performance of fast phylogenetic programs in the inference of single-gene trees. The bar-plots show the frequencies with which each of the six analysis strategies produced the best likelihoods for single-gene alignments in each of the (A) protein and (B) DNA data sets. Note that the best likelihood score for a given single-gene alignment can be found by more than one strategies; therefore the sum of frequencies for a data set may be greater than one.

Notably, the basic strategy *IQ-TREE* was the third best strategy with an overall frequency of 54.03%, slightly higher than that of the more comprehensive strategy *PhyML-10* (52.35%). In fact, *IQ-TREE* not only outperformed *PhyML-10* in 11 / 19 data sets, but also showed higher frequency than *RAxML-10* in the data set JarvD5b. On the other hand, *PhyML-10* performed consistently better than *RAxML* and *PhyML*, two basic strategies which were fifth and sixth, respectively, and had considerably lower overall frequencies (35.98% and 24.17%). Among basic strategies, *RAxML* performed better than *IQ-TREE* on only four (MisoA2, StruA5, MisoD2a, and MisoD2b) data sets, yet neither of them performed well on these data sets. Both *IQ-TREE* and *RAxML* found higher likelihood scores more often than *PhyML* in all data sets except for JarvD5b in which *RAxML* had slightly lower frequency.

In comparison, the likelihood scores obtained by *FastTree* were much lower than those of the other six strategies; the program produced the highest likelihood scores in only 1.67% of all alignments. However, *FastTree* also had substantial advantages in computational speed compared to the others (see below). Since FastTree can initiate tree searches using distinct starting trees, we performed additional FastTree analyses for selected data sets, consisting of 100 tree searches for each alignment starting from 50 parsimony trees and 50 random trees. The results show that in the vast majority of cases *FastTree* still generated worse likelihood scores than the other strategies even after compensating for the differences in runtime (supplementary table S3).

To further investigate the relative performance of the strategies using RAxML, PhyML, and IQ-TREE, we carried out pairwise comparisons between the three comprehensive strategies (i.e. *RAxML-10*, *PhyML-10*, and *IQ-TREE-10*) and also between their corresponding basic strategies (i.e. *RAxML*, *PhyML*, and *IQ-TREE*) (supplementary fig. S1). The overall trend is the same as that observed in fig. 2; on most data sets, *IQ-TREE-10* found better likelihood scores more frequently than *RAxML-10* which, in turn, outperformed *PhyML-10*; the same is true for the basic strategies. Interestingly, the three programs showed much closer performance when multiple trees searches were conducted. For instance, compared with *RAxML*, *IQ-TREE* found trees with equally good likelihood scores on 32.67% of all alignments and better scores for additional 20.59% of all alignments; the frequencies changed to 60.44% and 3.20%, respectively, in the comparison between *IQ-TREE-10* and *RAxML-10*. Nonetheless, *IQ-TREE-10* and *RAxML-10* still showed considerable advantages over *PhyML-10*; cumulatively, they found higher likelihood scores for additional 28.76% and 30.18%, respectively, of alignments than *PhyML-10*.

#### Tree topology

Trees with similar likelihood scores may differ substantially in their topologies, or *vice versa*. Hence, it is important to also examine the topological similarities between trees inferred by different methods in addition to their likelihood scores. Our evaluation is based on empirical data sets for which the true evolutionary histories are unknown, thus preventing a direct measurement of topological accuracy. Instead, we compared the trees inferred by various methods against the best-observed tree (i.e. the tree with the highest likelihood score) for each alignment. The rationale for using the best-observed ML trees as the references in our comparison is that, under the ML optimality criterion (which underlies all the methods examined here), the topologies of the trees with the highest likelihood scores are considered the best (currently known) answer.

We measured the normalized Robinson-Foulds, or nRF, distances (Robinson and Foulds 1981) between trees inferred by the seven strategies on each alignment against the corresponding best-observed tree. Overall, there was a strong positive correlation between the differences in likelihood scores and the topological distances when comparing inferred trees to the best-observed trees (Spearman’s correlations of 0.87 for all alignments and above 0.90 for most data sets, *p*-values < 2.2×10^-16^ in all cases). In other words, strategies that yielded likelihood scores closest or equal to the best-observed likelihood scores tended to be those whose topologies were also closest or identical to the best-observed topologies (supplementary table S4; see fig. 3 for data set YangA8 as an example).

**Figure 3.**
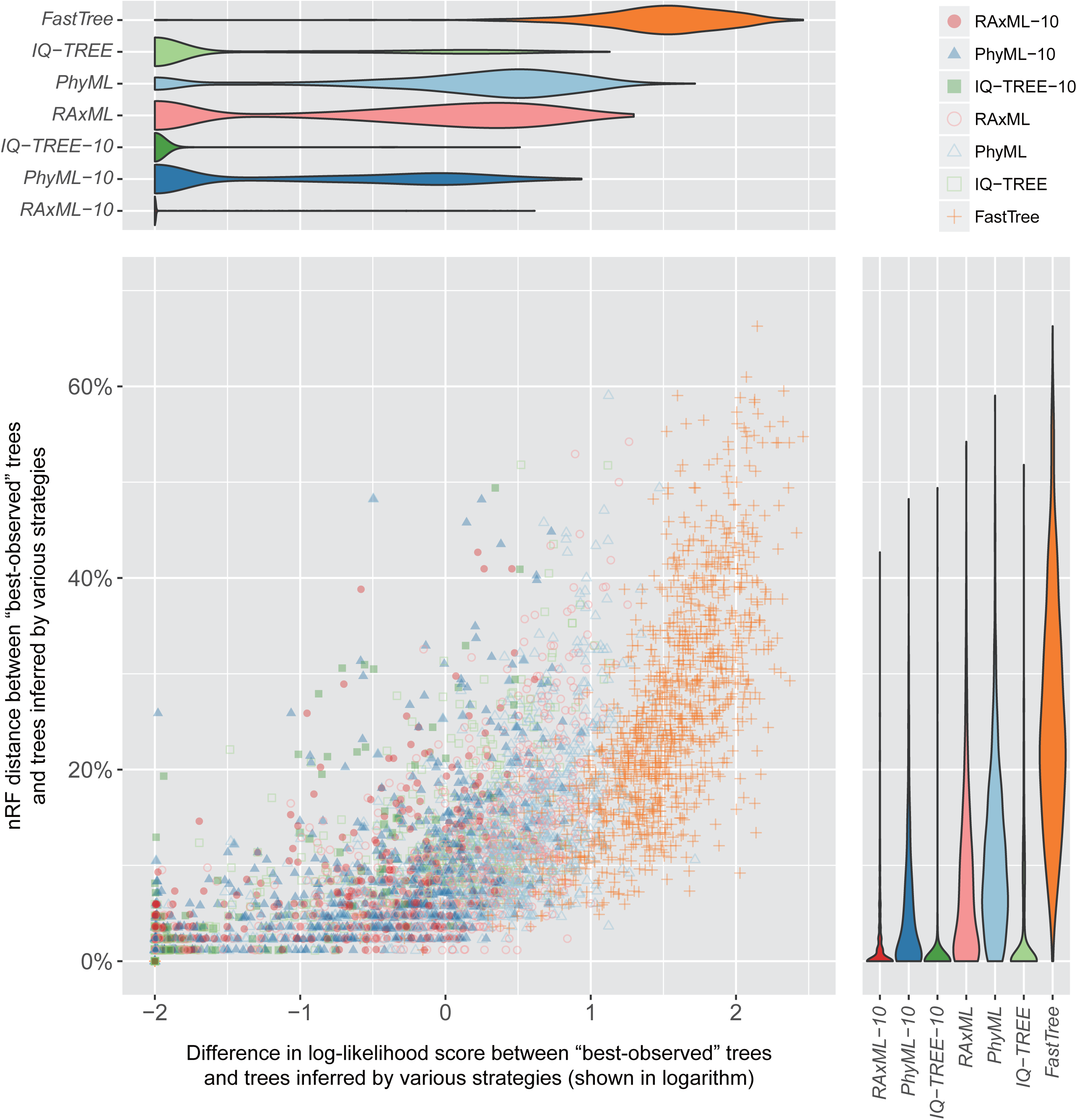
The performances of fast phylogenetic programs with respect to likelihood maximization and tree topology are positively correlated. Dots in the scatter plot correspond to trees inferred by various analysis strategies from single-gene alignments in data set A8. Log-likelihood score differences between inferred trees and the “best-observed” trees are plotted against the corresponding topological distances. The log-likelihood score differences are shown in logarithmic scale (with the addition of a small value of 0.01). The violin plots on the top and right show the distributions of log-likelihood differences (top) and topological distances (right), respectively, for trees inferred by each strategy.

Among the seven strategies, *IQ-TREE-10*, *RAxML-10*, and *IQ-TREE* showed the best performance in tree topology with median nRF distances of 0 for more than half of the data sets (supplemental table S4); this was unsurprising since these strategies contributed most of the best-observed trees. *PhyML-10*, *RAxML*, and *PhyML* also performed relatively well, with median nRF distances less than 0.03, 0.06, and 0.13, respectively, for most data sets. Here again, *FastTree* was behind the other strategies as it led to median nRF distances greater than 0.33 for most data sets.

#### Computational speed

To compare the computational speed of the seven strategies, we first measured the runtimes of *RAxML* (using a parsimony starting tree), *PhyML* (using a parsimony starting tree), *IQ-TREE*, and *FastTree*, as well as of RAxML and PhyML analyses using one random starting tree (referred to as *RAxML(RT)* and *PhyML(RT)*, respectively). We then plotted the runtimes of all these strategies against that of *RAxML* (fig. 4; supplementary table S5), and found strong positive correlations between the speeds of strategies over a wide range of runtimes (Spearman’ correlation ≥ 0.91 for all combinations of data types and strategies, *p*-values < 2.2×10^-16^ in all cases). The runtimes of *RAxML(RT)* and *PhyML(RT)* were highly similar to those of *RAxML* and *PhyML*, suggesting that *RAxML-10* and *PhyML-10* would take about ten times longer than *RAxML* and *PhyML*, respectively (supplementary table S6). Interestingly, *PhyML* was ∼1.5 times faster than *RAxML* on protein alignments, but ∼3.1 times slower on DNA alignments. On the contrary, *IQ-TREE* was faster than *RAxML* for both protein and DNA data (∼1.6 and ∼1.3 times faster, respectively), and the runtime of *IQ-TREE-10* would simply be ten times longer since it consists of ten independent IQ-TREE analyses. Lastly, *FastTree* was substantially more time-efficient than *RAxML* on both DNA alignments (∼47.9 times faster) and protein alignments (∼95.4 times faster). In addition, the time advantage of *FastTree* was greater for alignments requiring longer runtimes; for instance, our linear regression analysis suggests that *FastTree* might run ∼162.0 times faster than *RAxML* on the largest single protein alignments but only ∼9.6 times faster on the smallest ones.

**Figure 4.**
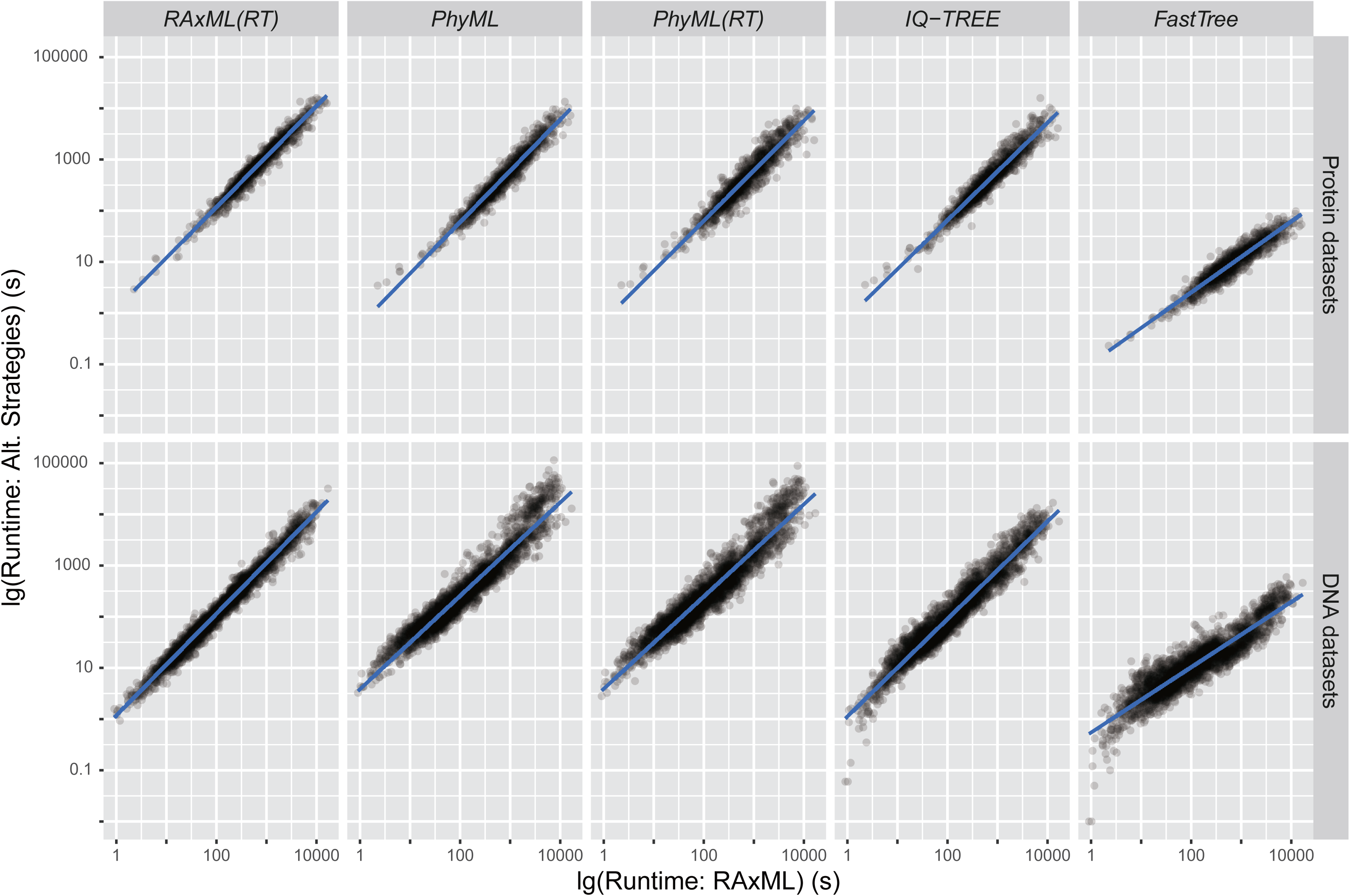
Runtime comparisons of fast phylogenetic programs in single-gene tree inferences. The runtimes required by each strategy to analyze a randomly selected subset of all protein (top row) and DNA (bottom row) alignments are plotted against the corresponding runtimes of *RAxML*. All runtimes (in seconds) are shown in logarithmic scale.

Overall, our results at the level of single-gene tree inference are consistent with previous, smaller-scale studies on the better efficiency of IQ-TREE relative to RAxML and PhyML (all using one search per alignment) (Nguyen et al. 2015), and the inferior performance of FastTree in likelihood score maximization as compared to other programs (Guindon et al. 2010; Liu et al. 2011). However, in contrast to previous observations (Guindon et al. 2010), we found that RAxML consistently outperformed PhyML in all data sets. This difference might be due to the small number of alignments examined in the previous study (Guindon et al. 2010) and the numerous updates of both programs since then.

#### Implications for efficient tree search on single-gene alignments

The inclusion of *RAxML-10*, *PhyML-10*, and *IQ-TREE-10* in our evaluation provided an opportunity to examine the effect of running multiple independent tree searches. For each of the three strategies, we first determined the highest likelihood score for each alignment, and then calculated the percentages of alignments for which the highest scores were found by given numbers of tree searches (supplementary fig. S2). In *IQ-TREE-10*, the highest likelihood scores were found in the first tree search for more than 70% of the alignments in 11 / 19 data sets (which explains the excellent performance of *IQ-TREE* in fig. 2), and the frequencies quickly approached 100% with additional tree searches. In contrast, the first tree search in *PhyML-10* found the highest likelihood scores for much fewer alignments (less than 30% in 10 / 19 data sets), and the frequencies increased more evenly with increasing numbers of tree searches. The plots of *RAxML-10* lie in between those of *IQ-TREE-10* and *PhyML-10* in most data sets. Interestingly however, in some data sets (e.g. MisoA2, StruA5, MisoD2a, MisoD2b), all three strategies showed almost the same linear increases in their frequencies of finding the highest scores with the number of tree searches (about 10% of the highest likelihood scores were found in each tree search). These results suggest that efficient tree search strategies are likely to vary between data sets and fast phylogenetic programs. To avoid unnecessary (or insufficient) tree search efforts, it is important to monitor the likelihood improvements over rounds of independent searches.

Additionally, the use of both parsimony and random starting trees in *RAxML-10* and *PhyML-10* allowed us to investigate the relative performance of the two types of starting tree. In our comparisons, parsimony and random starting trees showed comparable overall performance (supplementary fig. S3). For RAxML, five (or one) searches per alignment using random starting trees found better likelihood scores than using parsimony starting trees for only additional 3.47% (or 1.86%) of all alignments. In addition, equally good likelihood scores were obtained using both types of starting trees on 50.12% (or 31.73%) of all alignments when five (or one) RAxML searches were conducted. The same trend was also observed for PhyML. Together with their similar run-time performances (fig. 4), these results suggest that the two types of starting trees might be equally efficient in the analysis of single-gene alignments with moderate sequence numbers.

### Performance Test II: Coalescent-based species tree inference

In the second test, we assessed the fast ML-based phylogenetic programs in the context of the “two-step” coalescent-based species tree inference, in which single-gene trees were first estimated from individual alignments by each examined strategy and then used collectively to infer the species tree under a coalescent model (fig. 1A) (Liu et al. 2015). Here, we used the single-gene trees produced in the Performance Test I as input for the ASTRAL program (Mirarab and Warnow 2015), which was used to infer coalescent-based species trees. The species tree inferences by the seven strategies were then compared with the species tree estimated from the best-observed gene trees (referred to as best-observed species trees hereafter) to measure the topological distances (i.e. nRF distances).

We first determined for each data set the topological distances between the species tree inferred from the best-observed single-gene trees and those inferred from the gene trees inferred by each of the seven strategies. In that regard, the species tree estimations of all six strategies using RAxML, PhyML or IQ-TREE displayed comparably small topological distances to the best-observed species trees (median nRF distances ranged between0 and 0.03 across data sets), whereas the species trees inferred by *FastTree* were considerably more dissimilar (median nRF distances of 0.121) (table 3). When we only considered the bipartitions or splits that were strongly supported (i.e. had quartet-based posterior probability, or PP, support greater or equal to 0.9 [Sayyari and Mirarab 2016]), the species tree inferred by these strategies became even more similar to the best-observed species trees, although *FastTree*-generated species trees still showed the greatest topological distances (supplementary table S7). Nonetheless, for most strategies and data sets, the species tree estimates were much more similar to the best-observed trees than the corresponding single-gene tree inferences (table 3; supplementary table S8).

**Table 3.**
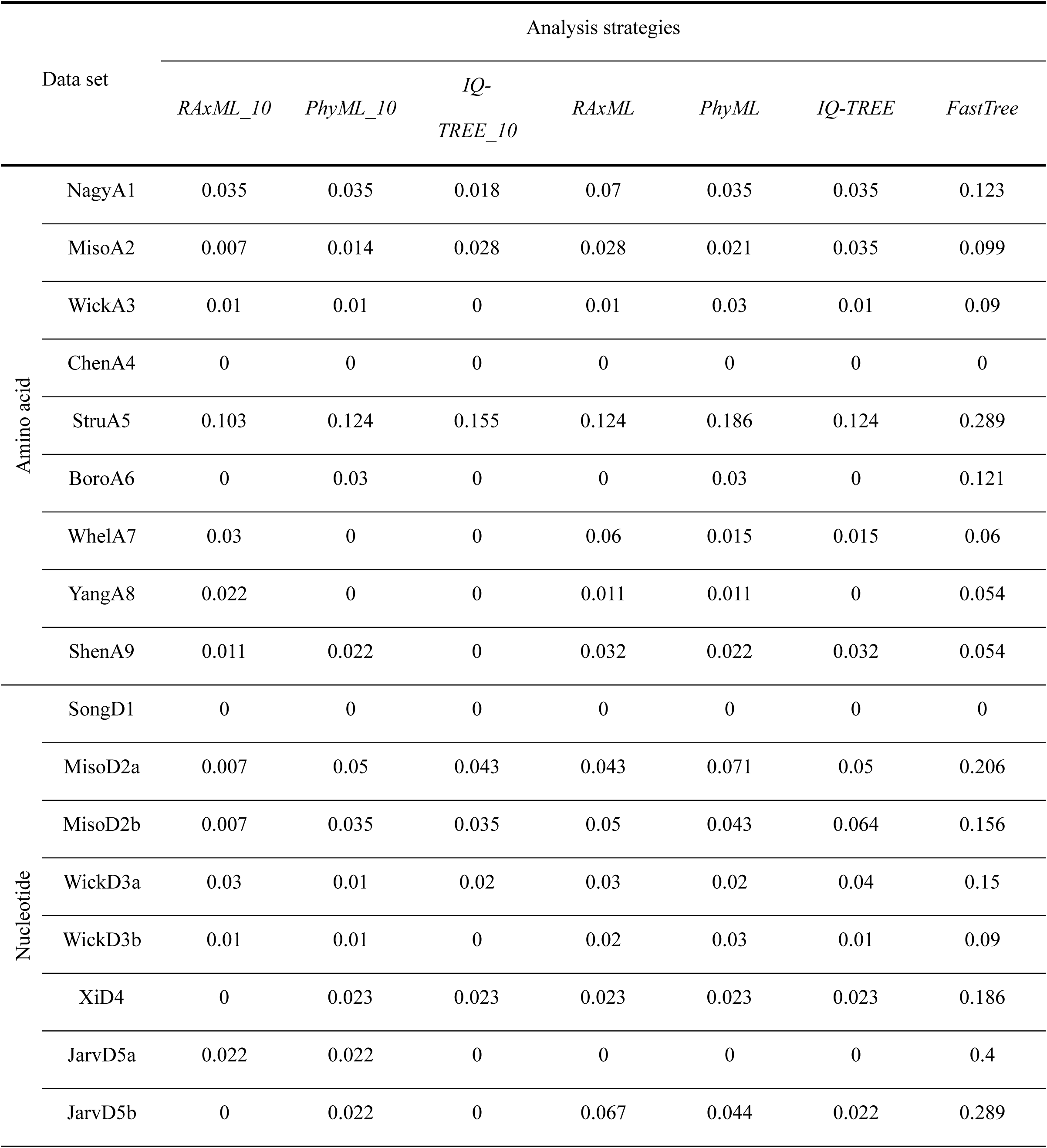
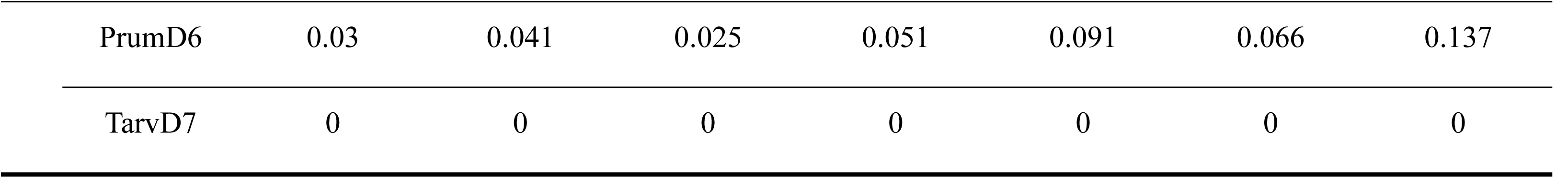
Normalized Robinson-Foulds distances between the coalescent-based species trees estimated from gene trees inferred by various strategies and the “best-observed” gene trees.

| Data set |  | Analysis strategies |  |  |  |  |  |  |
| --- | --- | --- | --- | --- | --- | --- | --- | --- |
|  |  | *RAxML_10* | *PhyML_10* | *IQ-*<br>*TREE_10* | *RAxML* | *PhyML* | *IQ-TREE* | *FastTree* |
| Amino acid | NagyA1 | 0.035 | 0.035 | 0.018 | 0.07 | 0.035 | 0.035 | 0.123 |
|  | MisoA2 | 0.007 | 0.014 | 0.028 | 0.028 | 0.021 | 0.035 | 0.099 |
|  | WickA3 | 0.01 | 0.01 | 0 | 0.01 | 0.03 | 0.01 | 0.09 |
|  | ChenA4 | 0 | 0 | 0 | 0 | 0 | 0 | 0 |
|  | StruA5 | 0.103 | 0.124 | 0.155 | 0.124 | 0.186 | 0.124 | 0.289 |
|  | BoroA6 | 0 | 0.03 | 0 | 0 | 0.03 | 0 | 0.121 |
|  | WhelA7 | 0.03 | 0 | 0 | 0.06 | 0.015 | 0.015 | 0.06 |
|  | YangA8 | 0.022 | 0 | 0 | 0.011 | 0.011 | 0 | 0.054 |
|  | ShenA9 | 0.011 | 0.022 | 0 | 0.032 | 0.022 | 0.032 | 0.054 |
| Nucleotide | SongD1 | 0 | 0 | 0 | 0 | 0 | 0 | 0 |
|  | MisoD2a | 0.007 | 0.05 | 0.043 | 0.043 | 0.071 | 0.05 | 0.206 |
|  | MisoD2b | 0.007 | 0.035 | 0.035 | 0.05 | 0.043 | 0.064 | 0.156 |
|  | WickD3a | 0.03 | 0.01 | 0.02 | 0.03 | 0.02 | 0.04 | 0.15 |
|  | WickD3b | 0.01 | 0.01 | 0 | 0.02 | 0.03 | 0.01 | 0.09 |
|  | XiD4 | 0 | 0.023 | 0.023 | 0.023 | 0.023 | 0.023 | 0.186 |
|  | JarvD5a | 0.022 | 0.022 | 0 | 0 | 0 | 0 | 0.4 |
|  | JarvD5b | 0 | 0.022 | 0 | 0.067 | 0.044 | 0.022 | 0.289 |

|  |  |  |  |  |  |  |  |
| --- | --- | --- | --- | --- | --- | --- | --- |
| PrumD6 | 0.03 | 0.041 | 0.025 | 0.051 | 0.091 | 0.066 | 0.137 |
| TarvD7 | 0 | 0 | 0 | 0 | 0 | 0 | 0 |

We further assessed the confidence levels (i.e. PP supports) of the incongruent bipartitions or splits identified in the abovementioned species tree comparison. Worryingly, the incongruent splits between the species tree inferred using *FastTree*-generated gene trees as input and the best-observed species tree received significantly higher PP supports (fig. 5; see supplementary table S9 for the results of Wilcoxon rank-sum tests); the median PP values of which were 0.81 for protein data sets and close to 1 for DNA data sets. Both of these values were much higher than those of the other six strategies, which were all below 0.60 and 0.71 for protein and DNA data sets, respectively.

**Figure 5.**
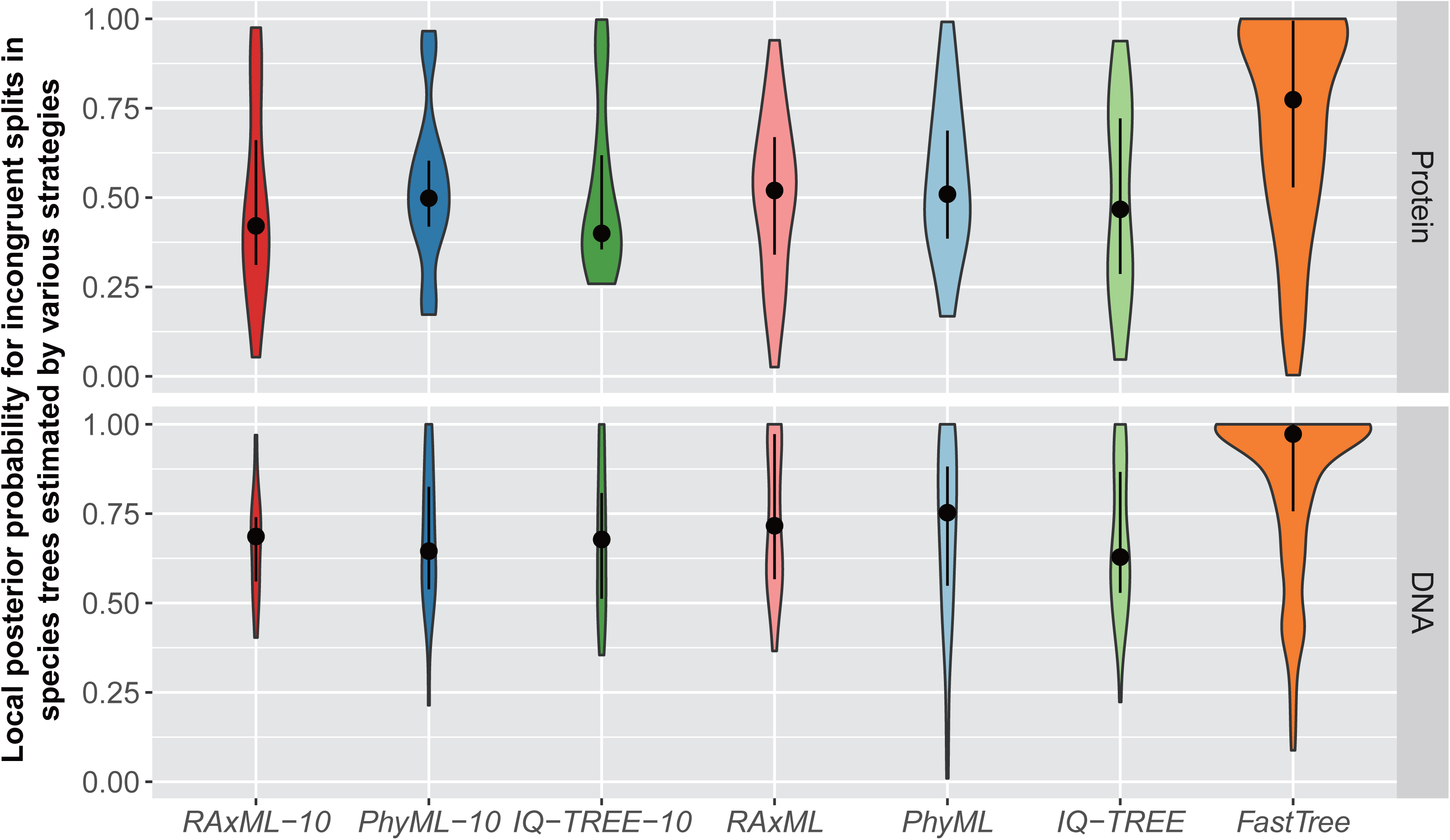
Incongruent splits in coalescent-based species trees estimated by the strategies using RAxML, PhyML, and IQ-TREE are weakly supported. The violin plots show the distribution of local posterior probabilities for incongruent splits in coalescent-based species trees estimated by various analysis strategies. Here, incongruent splits are defined as the splits that are not present in species trees estimated from best-observed single-gene trees. The areas of violin plots are proportional to the total numbers of incongruent splits. The grey dots and bars in each violin plot indicate the median and the first/third quartiles of the local posterior probabilities, respectively.

### Performance Test III: Concatenation-based species tree inference

In the third test, we examined the relative performance of the four programs in concatenation analysis of 17 taxon-rich and gene-rich supermatrices (we conducted concatenation analyses on 17, rather than 19, data matrices because: a) JarvD5a and JarvD5b correspond to different partitioning strategies from the same supermatrix [Jarvis et al. 2014], and b) MisoD2a does not have a corresponding supermatrix available from the original study [Misof et al. 2014]) (fig. 1B; table 2). Here, we again focused on the programs’ performance on likelihood score maximization, tree topology, and computational speed. However, as PhyML required exceedingly high runtime, memory, or crashed on multiple data sets, its results are not included in the evaluation. In addition to our analyses, all the supermatrices have also been previously extensively analyzed using either RAxML or ExaML (e.g. Jarvis et al. 2014; Misof et al. 2014; Wickett et al. 2014). Therefore, we included the reported likelihood scores and topologies – we refer to them as “RAxML/ExaML-published” trees – in our examination of relative performance.

#### Likelihood score maximization

Consistent with the pattern observed in single-gene tree analyses, RAxML and IQ-TREE achieved substantially higher likelihood scores than FastTree on supermatrix analyses (fig. 6; supplementary table S10). Interestingly, IQ-TREE found the highest likelihood scores in all 17 data sets and outperformed both our RAxML and previous RAxML/ExaML-published results on 7 and 8 data sets, respectively. Remarkably, IQ-TREE consistently yielded the highest likelihood scores in all independent replicates (except for the analyses of data set MisoD2a), while RAxML replicates were often trapped at suboptimal solutions (supplementary table S11). Moreover, the highest likelihood scores were usually found quite early in the IQ-TREE analyses (supplementary table S11), further suggesting its high efficiency in concatenation analysis.

**Figure 6.**
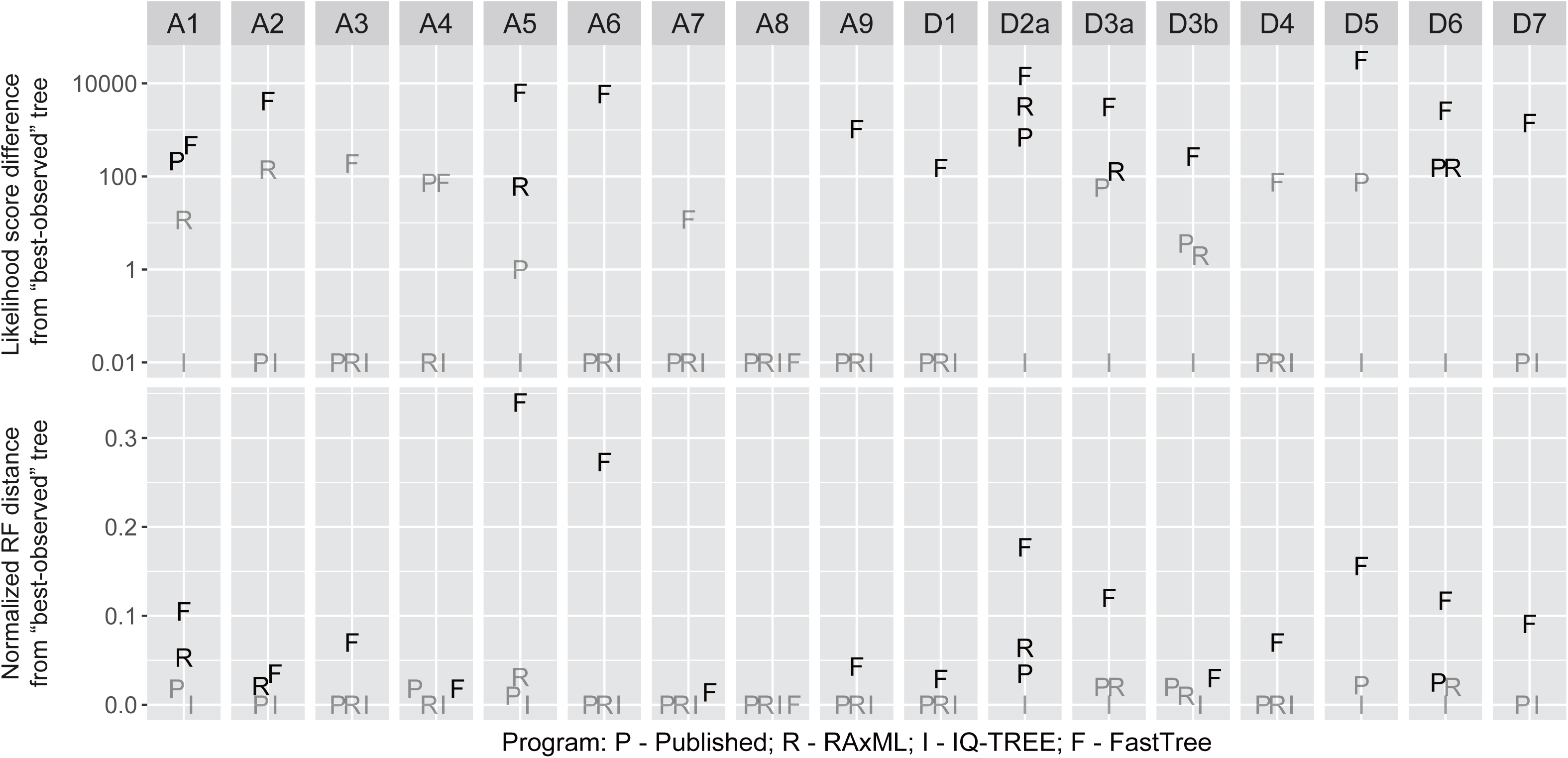
Likelihood score differences (A) and normalized Robinson-Foulds distances (B) between concatenation-based species trees inferred by various fast phylogenetic programs and the best-observed trees. In panel (A), the log-likelihood score differences are shown in logarithmic scale (with the addition of a small value of 0.01), and the likelihood scores that are not significantly different from the best-observed scores are shown in grey. In panel (B), the nRF distances of ExaML/RAxML-published and RAxML-generated trees that can be further improved by NNI rearrangements are shown in grey.

In comparison, RAxML/ExaML did not yield the highest likelihood scores for several data sets (fig. 6; supplementary table S10). One possible explanation is that, due to its “lazy SPR” heuristic, RAxML might report trees that are not optimal in terms of strict NNI or SPR rearrangement (Stamatakis 2015). Indeed, the best ML trees can be recovered by simply re-optimizing the RAxML-generated (or RAxML/ExaML-published) results using a function built in RAxML itself for four (or six) data sets (fig. 6; supplementary table S10). In addition, many of the differences in likelihood scores between trees inferred by RAxML/ExaML and IQ-TREE (the best ML trees) were small; three and five of the RAxML and previously published trees, respectively, were found to be equally good as the corresponding IQ-TREE trees as determined by approximately unbiased (AU) tests (fig. 6; supplementary table S10) (Shimodaira 2002). After taking these two factors into account, the likelihood scores of only one of our RAxML-generated trees and of two RAxML/ExaML-published trees that were significantly worse than their corresponding IQ-TREE results. In contrast, FastTree yielded significantly, and sometimes substantially, worse likelihood scores for most data sets. Furthermore, FastTree obtained lower likelihood scores than ExaML and IQ-TREE, even when it was allowed to run multiple times from distinct starting trees (supplementary table S12).

#### Tree topology

For all data sets, we calculated the nRF distances between trees inferred by the three programs and the best ML trees. As shown in figure 6, the topological distances of the examined programs are in agreement with their performance in likelihood score maximization (see also supplementary table S10). RAxML-generated or RAxML/ExaML-published trees were identical or highly similar to the best ML trees, with the largest nRF distance being 0.064. Importantly, some of the differences between the results of RAxML/ExaML and IQ-TREE correspond to contentious relationships in phylogenomic studies (e.g. in data set A4: the relative positions of pigeon, falcon, and other Neoaves; and in data set WickD3a: the relationships between Chloranthales, Eudicots, and Magnoliids) (Shen et al. 2017). Furthermore, some of these differences disappeared (and nRF distances became smaller) after the NNI-based re-optimization of RAxML/ExaML results. FastTree trees, on the other hand, showed much greater nRF distances from the best trees. We also evaluated the confidence levels (measures by Shimodaira–Hasegawa approximate likelihood ratio test, or SH-aLRT support [Guindon et al. 2010]) of trees that were significantly worse than the best ML trees. Figure 7 shows that large proportions of the incongruent splits in FastTree trees were highly supported.

**Figure 7.**
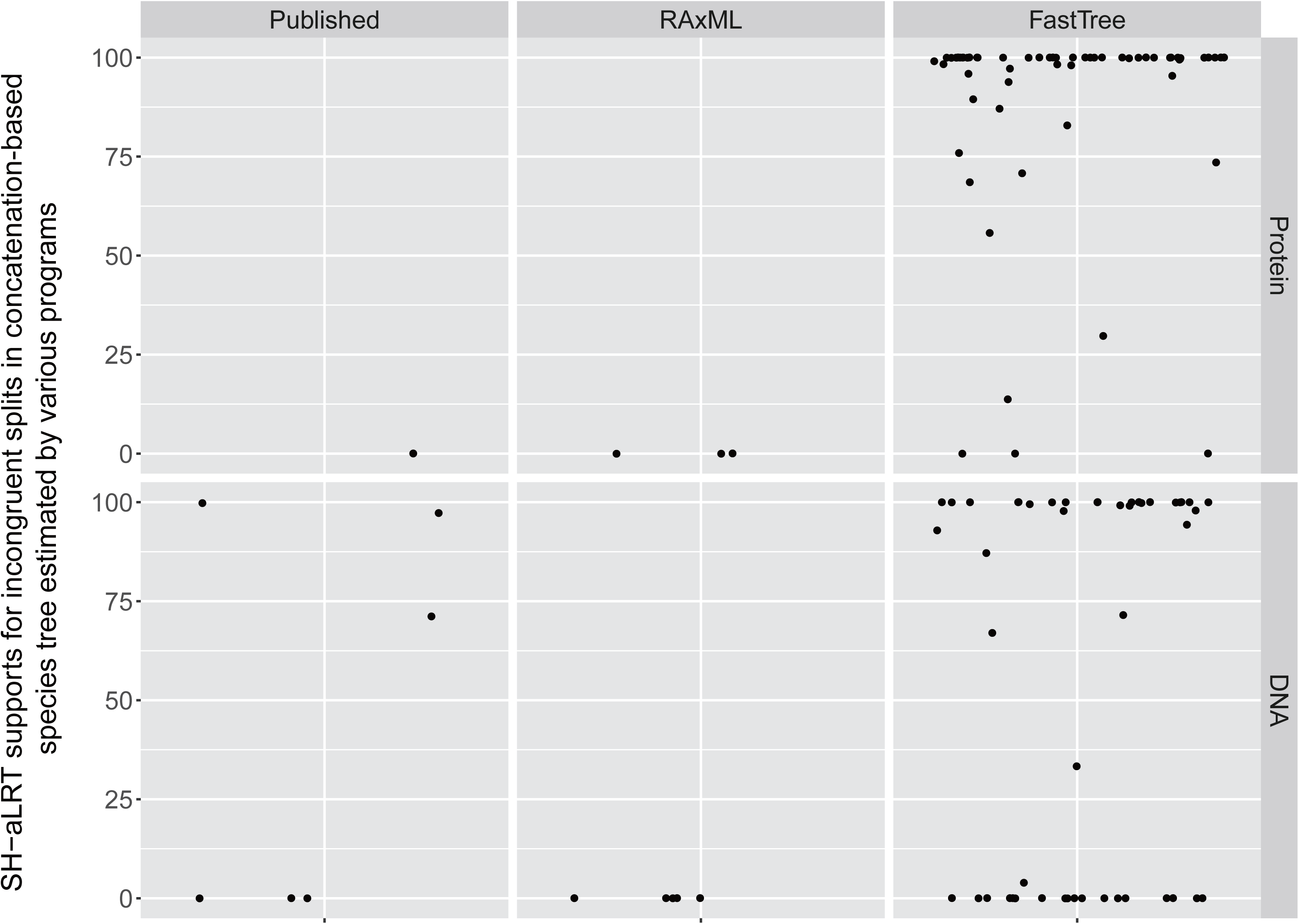
Many incongruent splits in concatenation-based species trees estimated by FastTree receive strong support. The jitter plots show the distribution of SH-aLRT supports for incongruent splits in concatenation-based species trees estimated by various fast phylogenetic programs. Here, incongruent splits are defined as the splits that are not present in the species trees with the best likelihoods. The species trees inferred by IQ-TREE contain no incongruent splits and therefore the data for IQ-TREE is not shown.

#### Computational speed

We compared the runtimes of ExaML, IQ-TREE, and FastTree on ten selected supermatrix data sets; each program was used to analyze each data set three independent times. The results are summarized in figure 8 (see also supplementary table S13). Overall, FastTree was significantly and substantially faster than ExaML and IQ-TREE (Wilcoxon signed-rank test, *p*-values < 0.01 for all pairwise comparisons), whereas the last two programs were on par with each other with respect to speed (Wilcoxon signed-rank test, *p*-value=0.56). Interestingly, IQ-TREE was faster on five of the six protein data sets, while ExaML was faster on all four DNA supermatrices.

**Figure 8.**
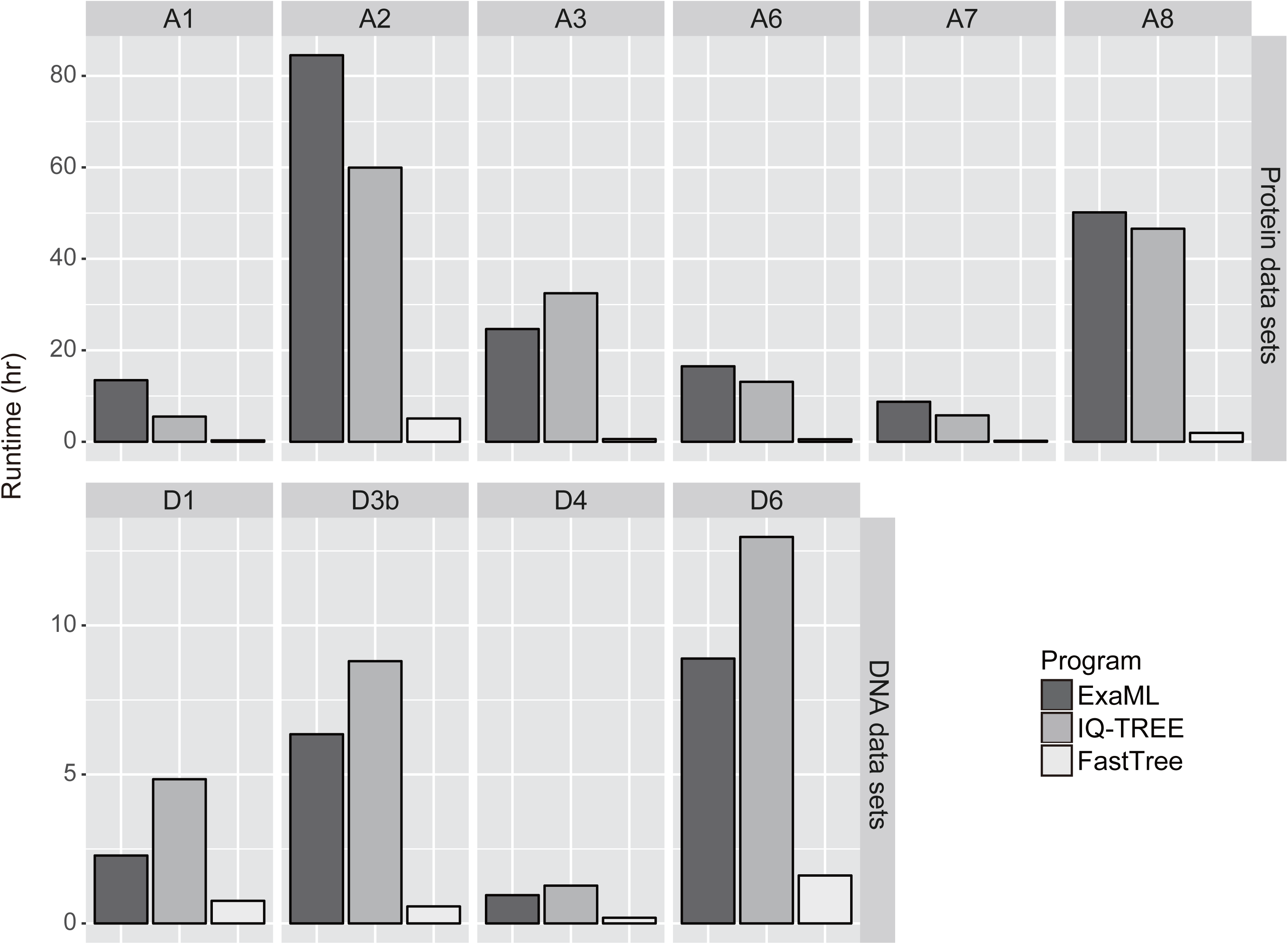
Runtime comparisons of fast phylogenetic programs in concatenation-based species tree inferences. The bar-plots show the runtimes (averaged over three replicates) required by RAxML, IQ-TREE, and FastTree to analyze 10 selected supermatrices.

These results suggest that IQ-TREE is a very appealing alternative to RAxML/ExaML, which is currently the default choice in most concatenation-based phylogenomic studies. This finding might not be entirely surprising because IQ-TREE represents the latest development in fast phylogenetic programs and has implemented a novel data structure to facilitate concatenation analysis (Chernomor et al. 2016). For RAxML and ExaML, our findings indicate that their results, even after multiple independent searches, should not be directly taken as the best answers and instead should be checked for potential improvements. On the other hand, together with the results of the coalescent-based test, our benchmarking suggests that FastTree is not suitable for production-level phylogenomic analyses. The exceptional runtime of FastTree might make it an attractive option for exploratory investigations, yet the results should still be interpreted with care.

### Impact of data properties on the relative performance of fast phylogenetic programs

In this benchmarking, we noticed several data properties that appear to have an influence on the relative performance of the examined programs. The first one is the number of sequences in the data set; in single-gene analyses, while IQ-TREE outperformed RAxML and PhyML in most instances, it did not do so on some of the data sets that had the largest numbers of taxa (MisoA2 and MisoD2a/b, 144 taxa; StruA5, 100 taxa; PrumD6, 200 taxa, when single tree search was performed; supplementary fig. S1). A potential explanation could be that IQ-TREE uses NNI as its topological rearrangement mechanism (Nguyen et al. 2015), whereas RAxML and PhyML are both based on SPR (Stamatakis 2006; Guindon et al. 2010). It is well recognized that SPR explores a greater proportion of tree space than NNI (Whelan and Morrison 2017) and that it does so in a manner proportional to the sequence number (SPR examines *O*(*n*^2^) neighbors for each tree instead of *O*(*n*) neighbors as by NNI). Therefore, while IQ-TREE exhibited better performance on data sets with fewer taxa through a combination of NNI rearrangement and stochastic algorithm, NNI might become a limiting factor on its performance on larger data sets.

Interestingly, in concatenation analyses, IQ-TREE found equally good or better trees than RAxML/ExaML for all data sets (fig. 6; supplementary table S10), including for the ones on which RAxML performed better in single-gene tree inference. The only difference between concatenation and single-gene tree analyses was the number of sites analyzed, a property that is strongly correlated with phylogenetic signal (Rokas et al. 2003; Shen et al. 2016a). Similarly, in single-gene tree inference, IQ-TREE showed much better performance over RAxML on data set JarvD5b than on JarvD5a (supplementary fig. S1); JarvD5b was derived from concatenating single-gene alignments in JarvD5a into a smaller number of longer partitions (Mirarab et al. 2014), resulting in enhanced phylogenetic signal (measured by average bootstrap support, or ABS, of gene tree) (supplementary fig. 4A). Compared with the single-gene data sets, these concatenated each tree instead of data matrices probably correspond to much simpler tree spaces in which the NNI algorithm might be sufficient. Consistent with this explanation, we found that the relative performance of SPR-based (RAxML and PhyML) and NNI-based (IQ-TREE) programs was indeed associated with the phylogenetic signal of alignment data. For instance, we compared the ABS values of the best-observed single-gene trees recovered by *RAxML-10* only, by *IQ-TREE-10* only, or by both programs, and found that they exhibited lower, intermediate, and higher ABS values, respectively (*p*-values < 2.2×10^-16^ for all Wilcoxon rank-sum tests; fig. 9). This trend held across most data sets (supplementary fig. 3A). We also observed the same pattern in the comparison between *PhyML-10* and *IQ-TREE-10* (supplementary fig. 4B), but not between *RAxML-10* and *PhyML-10* (supplementary fig. 4C). Investigating the relationship between the performance of fast phylogenetic programs and the strength of phylogenetic signal, which is in turn correlated with many other factors (Shen et al. 2016a), is an interesting area of future research.

**Figure 9.**
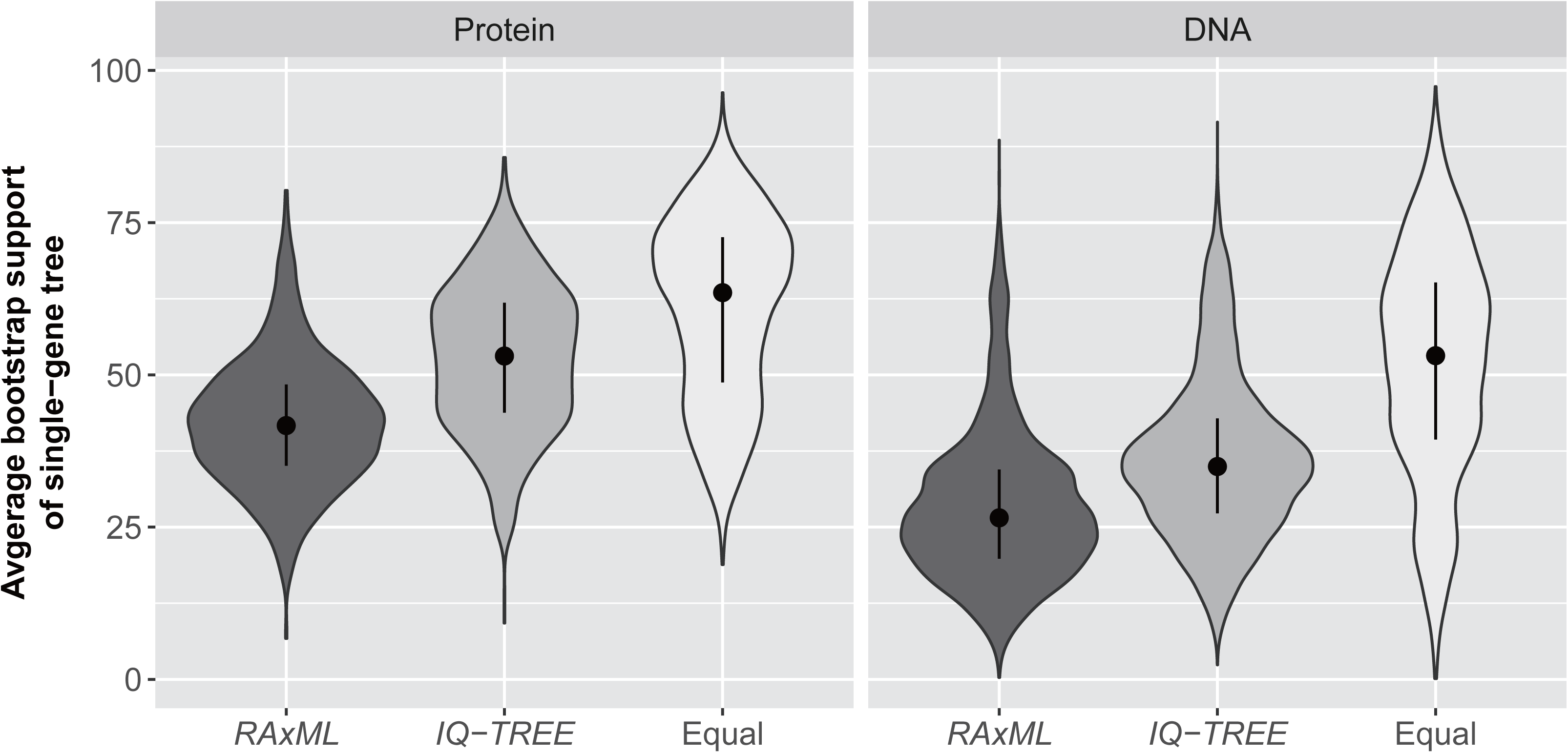
The strength of phylogenetic signal in the data has an impact on the relative performance of *RAxML-10* and *IQ-TREE-10*. The violin plots show the distributions of average bootstrap values of alignments for which the best likelihood scores were found by either *RAxML-10* or *IQ-TREE-10*, or both strategies at the same time. The average bootstrap values are taken from previously reported phylogenies for the alignments are used here as a measure of the strength of phylogenetic signal.

A hypothesis that stems from these results is that the performance of the SPR-based RAxML/ExaML programs will become more favorable (relative to that of the NNI-based programs) as the numbers of taxa included in phylogenomic data sets continue to increase beyond the numbers in the data sets examined in this study (i.e. 200 taxa in PrumD6). To that end, we further compared the performance of RAxML/ExaML and IQ-TREE on two supermatrices with much greater numbers of taxa, namely KatzA10 (800 taxa, 150 genes [Katz and Grant 2015]) and HugA11 (3083 taxa, 16 genes [Hug et al. 2016]) (supplementary table S1). Notably, all independent RAxML/ExaML searches were able to find better likelihood scores than IQ-TREE on both data sets (supplementary table S12), which is completely opposite to the results on data sets with 200 or less taxa. This result suggests that, in their current implementations, the SPR-based RAxML/ExaML is likely to be considerably more powerful than the NNI-based IQ-TREE in analyzing phylogenomic data sets that contain several hundreds or thousands of taxa.

Lastly, in agreement with previous studies (Guindon et al. 2010; Nguyen et al. 2015), we found that some programs displayed different time efficiency on protein and DNA data sets. For example, in single-gene analyses, PhyML was ∼1.5 times faster on protein alignments but ∼3.1 times slower on DNA alignments in comparison with RAxML (fig. 4; supplementary table S6). Similarly, in concatenation analyses, IQ-TREE required shorter runtimes than RAxML/ExaML on most protein data sets, while the opposite was true for DNA data sets. Such differential behavior may be attributed to the distinct algorithmic designs and/or software implementations of the programs on protein and DNA data (Guindon et al. 2010).

## Conclusion

In this study, we systematically examined and compared the performance of four popular, ML-based fast phylogenetic programs. As our evaluation covered standard phylogenetic and phylogenomic approaches (gene tree inference, as well as coalescent-based and concatenation-based species tree inference), assessed key parameters of inference (likelihood score, topology, and computational speed), and examined a comprehensive collection of empirical state-of-the-art phylogenomic data sets, our findings are directly relevant for the experimental design and execution of real-world phylogenetic and, particularly, phylogenomic studies.

## Materials and Methods

### Empirical phylogenomic data sets

The data sets were retrieved from their respective sources as listed in supplementary table S1. They were used in this study without any filtering on their contents, with two operations performed when necessary: 1) file split – some data sets (e.g. MisoD2) have only the concatenated alignments available, hence they needed to be split up to obtain single gene alignments; and 2) format conversion – alignments in the data sets are provided in either the “FASTA” or the “Phylip” formats, and had to be converted into the other format to be compatible with all examined phylogenetic programs (e.g. FastTree requires the “FASTA” format and PhyML requires the “Phylip” format). Similarly, all partition model files were transformed into the desired format for each phylogenetic program. Both the original and the actual files used for this study, as well as all the inferred trees are available from the figshare repository (https://figshare.com/account/home#/projects/22040, last accessed May 24, 2017).

### Single-gene tree inference

For single-gene tree inference, model selection analysis was first performed for each amino acid alignment to determine the best-fit model using the “TESTONLY” option of IQ-TREE v1.4.2 (Nguyen et al. 2015). The set of candidate models included all amino acid substitution models supported by RAxML, with and without empirical amino acid frequencies, and with the GAMMA correction for among site heterogeneity of evolutionary rates (Yang 1994) always enforced. For nucleotide alignments, the GTR model with empirical base frequencies and GAMMA distribution was used since it is the choice of almost all phylogenomic studies. Further details on the commands used for the model selection and all the analyses described below are available in the Supplementary Text.

Then each alignment was analyzed by single-threaded versions of the four fast phylogenetic programs. For the purpose of benchmarking, one tree search was conducted using each program under the same model settings (see below for FastTree as the only exception). We also performed additional RAxML searches with multiple parsimony and random starting trees, which represents a common strategy used in phylogenomic studies. In total, seven strategies of phylogenetic analysis were assessed:

1. *RAxML-10*: Two analyses were carried out for each alignment using RAxML v8.2.0 (Stamatakis 2014); one included five independent searches starting from parsimony trees and the other five starting from random trees. A random number seed was generated independently and fed into each analysis. The BFGS optimization method was turned off in the analyses of nucleotide alignments since it has been reported previously to produce unstable results (Church et al. 2015). The likelihood scores of the trees inferred by the two analyses were compared to determine the final result of *RAxML-10* and the tree with the highest likelihood was selected; in cases where two trees had equally high likelihood scores but different topologies, a random selection was made from the two trees (see the “Assessment of tree inferences” section for detailed procedure on likelihood score and topological distance calculations);
2. *RAxML*: One search was carried out for each alignment using RAxML v8.2.0 (Stamatakis 2014) with a parsimony starting tree. The analysis was initiated using the same random seed number as the analysis based on parsimony starting tree in *RAxML-10*, and thus can be considered as a subset of the tree inferences conducted in *RAxML-10*. Therefore, *RAxML-10* will always produce equal or better results than *RAxML*. All other settings were the same as *RAxML-10*;
3. *PhyML-10*: Five independent analyses were carried out for each alignment using PhyML v20160530 (Guindon et al. 2010); each included one search starting from a parsimony tree and one other search starting from a random tree. The “SPR” algorithm was selected for tree topology search. Certain amino acid substitution models (e.g. JTTDCMut and mtZOA) were specified as custom models since they were not supported by PhyML natively. Unlike in RAxML analyses, random number seeds were generated automatically by PhyML. The tree with the highest likelihood was selected in the same way as in *RAxML-10*;
4. *PhyML*: One single search on each alignment using PhyML v20160530 (Guindon et al. 2010) with a parsimony starting tree, corresponding to the first parsimony starting tree-based search in *PhyML-10*.
5. *IQ-TREE-10*: Ten independent searches were carried out for each alignment using IQ-TREE v1.4.2 with default settings except for the model. Similar to PhyML, IQ-TREE generates random seed numbers automatically. The tree with the highest likelihood was selected in the same way as in *RAxML-10*;
6. *IQ-TREE*: One search on each alignment using IQ-TREE v1.4.2, corresponding to the first tree search in *IQ-TREE-10*;
7. *FastTree*: One search was carried out for each alignment using FastTree v2.1.9 (Price et al. 2010) with the default heuristic NJ starting tree. The “-spr 4”, “-mlacc 2”, and “-slownni” options were specified to enable more thorough heuristic tree search. Unlike the other programs, FastTree only supports three amino acid substitution models (i.e. JTT, WAG, and LG). Therefore, the best-fit model among the three was selected for each FastTree analysis of amino acid alignment. Moreover, the algorithm of FastTree is deterministic, thus independent analyses of the same alignment will always lead to the same result.

Once all single-gene tree estimations were completed, each alignment was associated with at least seven gene trees, which included the trees inferred by the seven above-mentioned strategies and, for most data sets, previously reported single-gene trees from respective publications. The gene trees of each alignment were then compared to identify the one with the best likelihood score, which is referred to as the “*best-observed*” tree; the tree with the highest likelihood score was selected to be the best-observed tree, or, if multiple trees had the same likelihood score, a random selection was made among them (see the “Assessment of tree inferences” section for detailed procedure on likelihood score and topological distance calculations).

### Coalescent-based species tree inference

Each of the 19 data sets was analyzed following the “two-step” procedure of coalescent-based species tree inference (Liu et al. 2015); single-gene trees were first estimated using fast ML-based phylogenetic programs (see above) and were then used to infer the species tree with the coalescent-based approach implemented in the ASTRAL program, v4.10.12 (Mirarab and Warnow 2015). In total, eight coalescent-based species trees were estimated for each data set, seven of which were based on single-gene trees produced by the seven strategies, and the eighth one was based on the “best-observed” trees.

### Concatenation-based species tree inference

Supermatrices consisting of all single-gene alignments and corresponding model files indicating partition scheme as well as model assignments are available for all data sets except for MisoD2a and JarvD5b. Concatenation-based species tree inferences were performed on these supermatrices using parallelized versions of all phylogenetic tools whenever possible due to the heavy computation being required. Edge-linked partitioned analyses (i.e. branch-lengths shared across partitions) were performed on each supermatrix using both RAxML and IQ-TREE. The RAxML analyses were conducted using RAxML-MPI v8.2.3 (available through the CIPRES Scientific Gateway), each consisting of six to eight tree searches with parsimony starting trees, while five independent IQ-TREE searches were carried out for each supermatrix using IQ-TREE-OMP v1.4.2. FastTreeMP v2.1.9 was run once per supermatrix with the thorough search parameters (see above); partition schemes were not used since FastTree does not support partitioned analysis. PhyML v20160530 was also used to analyze the supermatrices but failed on multiple data sets (the analyses either collapsed or did not finish after more than one week of computation).

### Assessment of tree inferences

In order to evaluate the performance of different fast phylogenetic programs, their inferred trees were compared from the following three aspects:

1. Likelihood: With respect to likelihood score maximization, a program was considered to perform better than another if it yielded a log-likelihood score that was more than 0.01 higher than the other. To ensure the fairness of the comparison, the likelihood scores of all trees were re-calculated using RAxML v8.2.0 with models set to the best-fit models and “GTR+G” for amino acid and nucleotide single-gene alignments, respectively, or the respective partition schemes for supermatrices. Trees of the same topology are presumed to have the same likelihood score. The BFGS optimization method was turned off in the analyses of nucleotide alignments. Independent likelihood score re-calculations were conducted for all trees using IQ-TREE v.1.4.2 and the R package “phangorn” v2.2.0 to control for potential biases since RAxML itself is one of the programs to be assessed. The results were essentially the same (Spearman’s correlations ≥ 0.99 and *p*-values < 2.2×10^-16^ for all pairwise comparisons; supplementary fig. S5; supplementary tables S14 and S15);
2. Topology: Our benchmarking is based on empirical data sets whose true underlying histories were unknown, thus preventing a direct measurement of the topological accuracy of programs. Thus, we compared the trees inferred by various strategies/programs against the tree with the best likelihood score observed for each alignment by calculating the pairwise Robinson-Foulds (RF) distances (Robinson and Foulds 1981) between them. To allow for comparison across alignments, the RF distances were normalized by the total numbers of internodes in respective pairs of trees. The reliabilities of coalescent-based and concatenation-based species tree estimations were evaluated using the local posterior probability measure (Sayyari and Mirarab 2016) implemented in ASTRAL v4.10.12 and the SH-aLRT test (Guindon et al. 2010) implemented in IQ-TREE v1.4.2, respectively;
3. Speed: Computational efficiency is another critical factor affecting the choice of phylogenetic programs, especially when the availability of computational resource is a concern. The aforementioned phylogenetic analyses were conducted on multiple different computational platforms, each equipped with different types of CPUs, thus preventing a direct comparison of the runtimes. To address this issue, we selected 10% of single-gene alignments randomly from each data set and redid all relevant phylogenetic analyses on Vanderbilt University’s ACCRE cluster (http://www.accre.vanderbilt.edu/) using the same type of computing nodes. Similarly, a subset of supermatrices were selected and re-analyzed by ExaML v3.0.17 and IQ-TREE-OMP v1.4.2 (each with three replicates) on the same type of ACCRE nodes.

### Computational resources

In this study, we conducted more than 670,000 tree inferences on about 45,000 single-gene alignments and supermatrices, which costed more than 300,000 CPU hours of computational time in total. This huge amount of phylogenetic analyses was made possible by using three supercomputing resources, including the Advanced Computing Center for Research and Education (ACCRE) at the Vanderbilt University, the University of Wisconsin-Madison Center for High Throughput Computing (CHTC), and the CIPRES Scientific Gateway at the San Diego Supercomputer Center (Miller et al. 2010). Single-gene analyses were distributed between ACCRE and CHTC. For supermatrices, RAxML analyses were performed using the “RAxML-HPC v.8 on XSEDE” interface on CIPRES, while the other analyses were carried out on ACCRE.

## Acknowledgements

We thank Dr. Jacek Kominek for helpful suggestions. This work was conducted in part using the resources of the Advanced Computing Center for Research and Education at Vanderbilt University, the University of Wisconsin-Madison Center for High Throughput Computing, and the CIPRES Science Gateway at the San Diego Supercomputer Center. This work was supported by the National Science Foundation (DEB-1442113 to A.R.; DEB-1442148 to C.T.H.), in part by the DOE Great Lakes Bioenergy Research Center (DOE Office of Science BER DE-FC02-07ER64494), and the USDA National Institute of Food and Agriculture (Hatch project 1003258 to C.T.H.). C.T.H. is a Pew Scholar in the Biomedical Sciences, supported by the Pew Charitable Trusts.

